# Social anxiety links of amygdala switch cross puberty

**DOI:** 10.1101/2024.05.01.591960

**Authors:** Quan Zhou, Zi-Xuan Zhou, Yin-Shan Wang, Xi-Nian Zuo

## Abstract

Socioemotional functions have been linked to the amygdala through molecular mechanisms observed in animals. However, these links remain largely unestablished in the development of the human amygdala during puberty. By precisely tracing the amygdala with longitudinal data spanning childhood and adolescence, our aim is to capture the dynamic relationship between social anxiety and amygdala geometry. Around the onset of puberty (10-11.5 years), we detected shifting associations between amygdala volume and social anxiety. Higher social anxiety is associated with a larger amygdala in mid-childhood but a smaller amygdala in early adolescence. Further geometry analysis revealed regional deformations that underpinned the shift. Our findings reconcile inconsistent results from previous studies and respect the intrinsic dimension of development in resolving amygdala-anxiety links during puberty.

---

Group living offers significant adaptive advantages for numerous species, particularly primates, necessitating the evolution of complex social abilities (*1*). Adolescence represents a critical phase for the development of socioemotional functions, supported by pubertal processes that encourage adolescents to develop or refine socioemotional skills and networks in a context-sensitive way (*2, 3*). The onset of puberty activates hormonal changes, leading to significant brain development (*3–6*), which, in turn, increases the sensitivity of adolescents to social contexts and evaluations (*7, 8*). This period also sees an increased need for peer acceptance (*9*), which could raise social stress levels (*10, 11*). A previous study reported an increase in social anxiety and fear during childhood, peaking in adolescence (*12*). In particular, as a hub in brain networks that support social life (*13, 14*), the amygdala undergoes distinct neurodevelopmental changes related to social functioning throughout adolescence, and many neural networks in which the amygdala participates undergo functional reorganization to support emerging and changing behaviors in adolescence (*15–20*). However, the development of the association between amygdala geometry and social anxiety from childhood to adolescence is poorly understood. Previous research in children demonstrated a positive correlation between amygdala volume and social anxiety, showing that children with an anxious or inhibited temperament - a risk factor for social anxiety disorder - have a larger amygdala than their less inhibited counterparts (*21, 22*). Sparks et al. followed a large sample of 3- to 4-year-old children, finding that only individuals with more severe social impairments in children with autism showed amygdala enlargement compared to typically developing children (*23*). In contrast, a study in adolescents with an average age of 13 years enrolled adolescents with pediatric anxiety, particularly social phobia, who showed a smaller amygdala than healthy adolescents (*24*). These findings suggest inconsistent associations between amygdala structure and social anxiety during childhood and adolescence, with associations that possibly vary depending on the subject’s age (*25*). Although Suor et al. (*26*) have not found that age moderates the relationship between social anxiety and amygdala volume, limitations in sample size (small convenience samples, N=35) severely restrict the generalizability of the results, underscoring the need for more investigations from a developmental population perspective.

We aim to elucidate how associations between amygdala geometry features (i.e., volume and shape) and social anxiety evolve through childhood and adolescence. This would help inform a more complete understanding of the neural mechanisms that underlie the formation and development of social competence. We first established the relationship between social anxiety and amygdala volumes, obtained by precisely tracing 385 longitudinal structural magnetic resonance images of 185 healthy children and adolescents (baseline age: 7-16 years) with the gold standard (i.e., manually). The shape analysis was then used to model the geometry deformation of the amygdala and chart its time-varying relationship with social anxiety from childhood to adolescence.

## Results

### No detectable static link between social anxiety and amygdala volume

The regression analysis using the generalized additive mixed model (GAMM) identified the optimal models with sex as a fixed effect factor and anxiety adjusted for a smoothing spline function (see Table 1). This model indicated the redundancy of a fixed age effect and an interaction term between anxiety and sex, and revealed a larger amygdala in boys than in girls, but the changing rates of the regression curves did not differ by sex. As in Table 1 and Figure 1, no significant correlation was detected between social anxiety and bilateral amygdala volumes in girls and boys during childhood and adolescence (left amygdala: *F* =0.13, *p*=0.719; right amygdala: *F* =0.19, *p*=0.660). Furthermore, the influence of sex on the bilateral amygdala volume is significant, manifesting itself in significant sex differences at the intercept of the regression curve (left amygdala: *t* =5.09, *p* < 0.05; right amygdala: *t* =4.45, *p* < 0.05).

**Figure 1:**
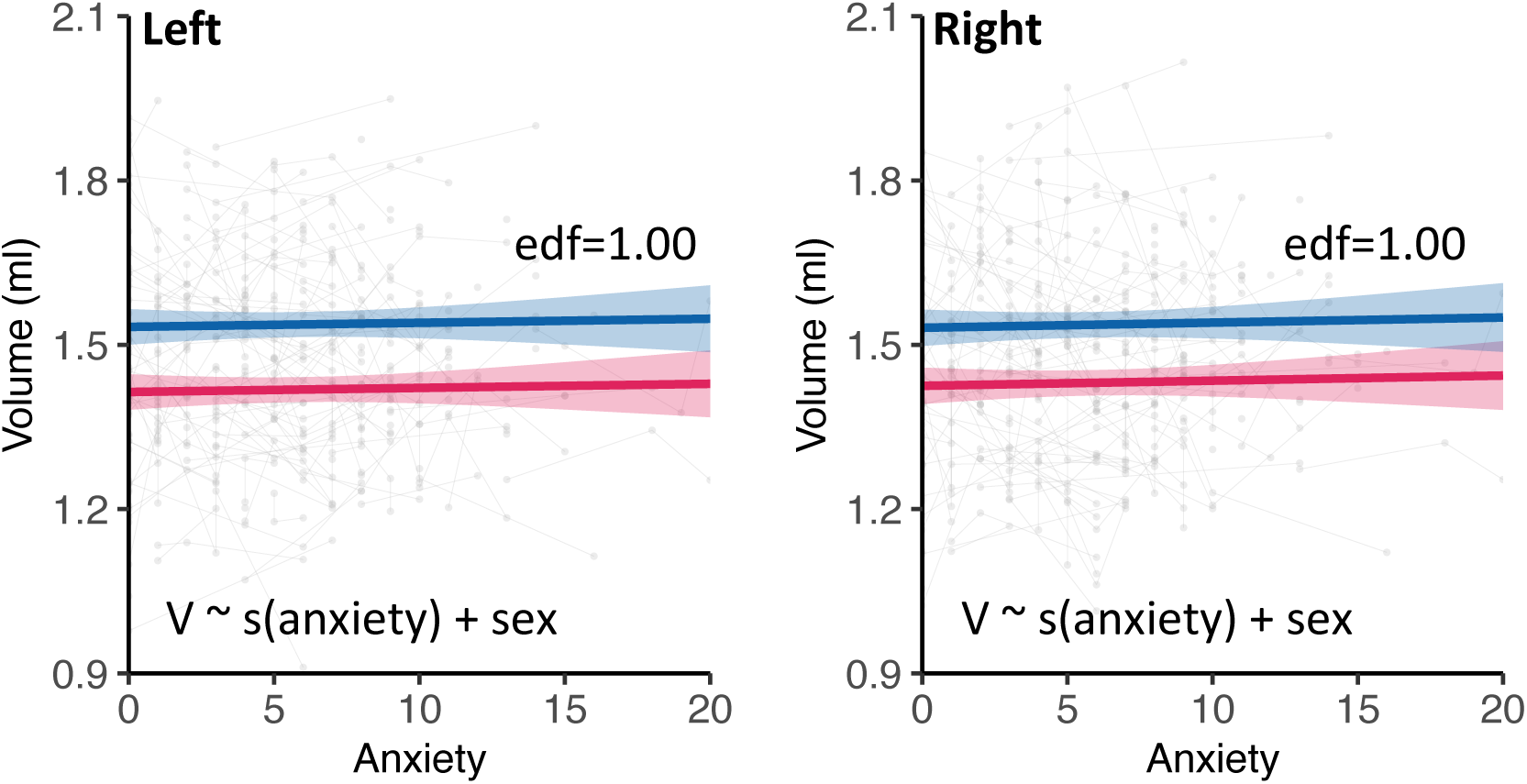
Associations between social anxiety and left/right amygdala volume. No significant correlation was detected between social anxiety and volumes of the left and right amygdala, as indicated by the dark lines and their surrounding shaded 95% confidence intervals.

**Table 1:** GAMM estimates of the social anxiety effect on amygdala volume (6-18 years)

| Hemi | Best | R<sup>2</sup> | Sex |  |  |  | Anxiety spline |  |  |  |
| --- | --- | --- | --- | --- | --- | --- | --- | --- | --- | --- |
|  | model fit | (adjusted) | Estimate | SE | *t* | *p* | edf | Red.df | *F* | *p* |
| Left | s(anx) + sex | 0.12 | 0.12 | 0.02 | 5.09 | <**0.001** | 1.00 | 1.00 | 0.13 | 0.719 |
| Right | s(anx) + sex | 0.09 | 0.11 | 0.02 | 4.45 | <**0.001** | 1.00 | 1.00 | 0.19 | 0.660 |

### Time-varying links between social anxiety and amygdala volume

The analyses of the time-varying effect model (TVEM) revealed that the relationship between social anxiety and amygdala volume changes at different stages of development (see Figures 2C and 2D). A positive correlation between social anxiety and bilateral amygdala volumes was observed during mid-childhood (see Figures 2C and 2D). However, this relationship changes in early adolescence, where social anxiety scores were found to negatively correlate with bilateral amygdala volumes (Figure 2C and 2D). Post hoc GAMMs revealed patterns in the correlations between social anxiety and bilateral amygdala volume in mid-childhood and early adolescence (Table 2). The analysis determined that the best fitting growth curves for each age stage, as indicated by the AIC, were those that employed smoothing anxiety models. For the bilateral amygdala, the second model was favored by the AIC for mid-childhood, and the first model was favored for early adolescence (Table 2). As shown in Figures 2A and 2B, bilateral amygdala volumes were significantly positively correlated with social anxiety at 8-10 years (left amygdala: F=5.94, p=0.017; right amygdala: F=5.69, p=0.011), indicating that larger volumes of amygdala were related to increased social anxiety in mid-childhood, while around 11.5-13 years (Figures 2E and 2F), bilateral volumes of amygdala were negatively correlated with social anxiety (left amygdala: F=5.06, p=0.027; right amygdala: F=4.21, p=0.016), indicating that smaller volumes of amygdala were related to increased anxiety in early adolescence. These findings collectively indicate a transition in the associations between social anxiety and amygdala volume from positive in mid-childhood to negative in early adolescence.

**Figure 2:**
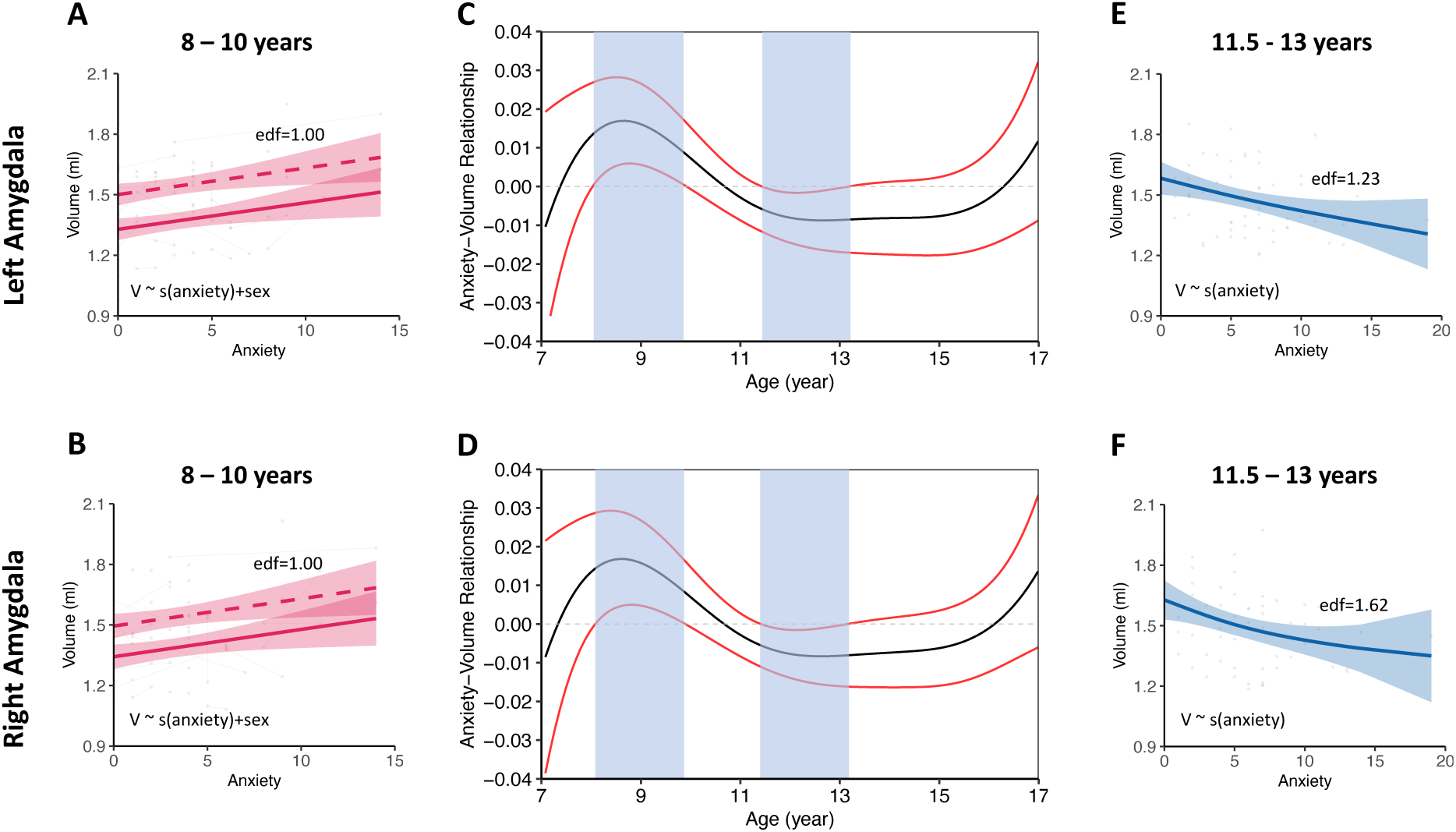
Effects of social anxiety on left (C) and right (D) amygdala volume across childhood and adolescence. For the bilateral amygdala, higher social anxiety is associated with larger amygdala volume in mid-childhood, whereas in early adolescence, it is associated with smaller amygdala volume. The black lines depict the trajectory of the association between social anxiety and amygdala volume over age, with the red lines representing the 95% confidence intervals. The transparent blue boxes highlight periods when social anxiety is significantly associated with amygdala volume. The plots illustrate the relationships between social anxiety and left amygdala volume in (A) mid-childhood (8-10 years) and (E) early adolescence (11.5–13 years), and between social anxiety and right amygdala volume in (B) mid-childhood (8-10 years) and (F) early adolescence (11.5–13 years).

**Table 2:** Post-hoc GAMM estimates of anxiety for bilateral amygdala volume.

| Age range | Best<br>model fit | N | R<sup>2</sup> | Sex |  |  |  | Anxiety spline |  |  |  |
| --- | --- | --- | --- | --- | --- | --- | --- | --- | --- | --- | --- |
|  |  |  |  | Estimate | SE | *t* | *p* | edf | Red.df | *F* | *p* |
| Left Hemi |  |  |  |  |  |  |  |  |  |  |  |
| 8.00-10.00 | s(anxiety) + sex | 78 | 0.24 | 0.17 | 0.04 | 4.06 | <**0.001** | 1.00 | 1.00 | 5.94 | **0.017** |
| 11.50-13.00 | s(anxiety) + sex | 78 | 0.16 | 0.15 | 0.05 | 3.17 | **0.002** | 1.00 | 1.00 | 5.06 | **0.027** |
| Right Hemi |  |  |  |  |  |  |  |  |  |  |  |
| 8.00-10.00 | s(anxiety) | 61 | 0.11 |  |  |  |  | 1.23 | 1.23 | 5.69 | **0.011** |
| 11.50-13.00 | s(anxiety) | 61 | 0.12 |  |  |  |  | 1.62 | 1.62 | 4.21 | **0.016** |

### Amygdala shape deformation underlying the dynamic anxiety links

To further explore the shape deformation involved in the observed dynamic links between social anxiety and amygdala volumes during development, we performed a post hoc analysis of the association of shape with social anxiety. Regarding the post hoc nature of this procedure and the robustness consideration (*27*), the statistical significance of the association tests remained uncorrected in the vertex for multiple comparisons. As shown in Figure 3, the distribution map of the *p* values revealed areas at the vertex level (”hot spots”) of anxiety-related areal development, a very small region located in the basal nucleus of the amygdala.

**Figure 3:**
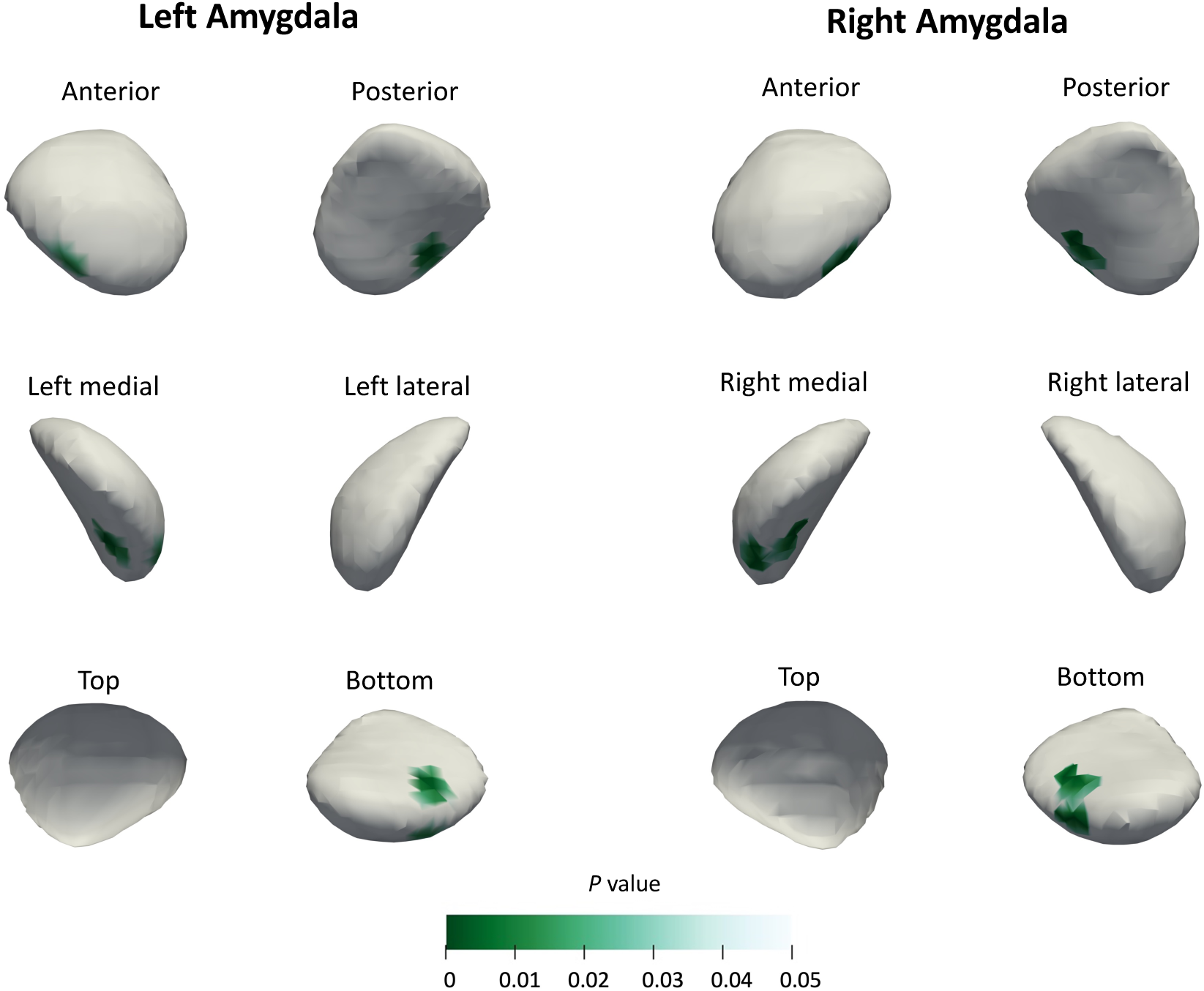
Statistically significant surface deformations associated with social anxiety. For the bilateral amygdala, the anterior, posterior, medial, lateral, top, and bottom view of *P*-statistic maps were shown. Significant surface vertices (0.05) are colored in green.

Figure 5 shows the two primary PCs (PC1 and PC2) for the bilateral amygdala, indicating that the high PC1 loading regions were primarily concentrated in the laterobasal subnuclei of the amygdala, while the low PC1 loading regions were mainly located in the superficial and centromedial subnuclei (variance explained by PC1: left amygdala: 41%; right amygdala: 50%). The spatial pattern of PC2 loading was different from that of PC1, with the high PC2 loading regions mainly concentrated in the centromedial and superficial subnuclei, while the low PC2 loading regions were concentrated in the lateral subnuclei of the laterobasal amygdala (variance explained by PC2: left amygdala: 29%; right amygdala: 23%).

**Figure 4:**
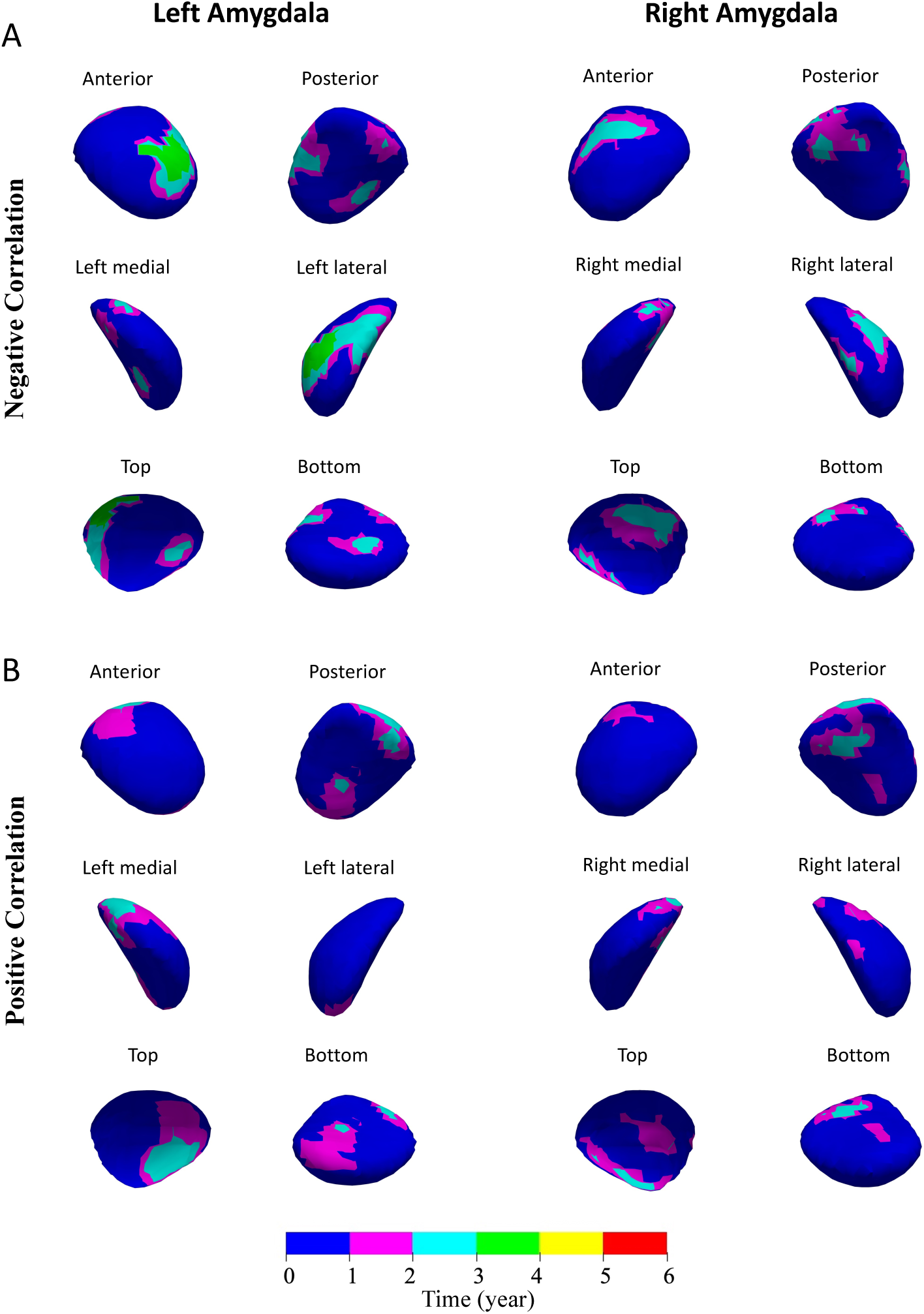
Total duration between 7 and 16 years of the statistically (A) negative and (B) positive relationship between the regional amygdala surface area and social anxiety.

**Figure 5:**
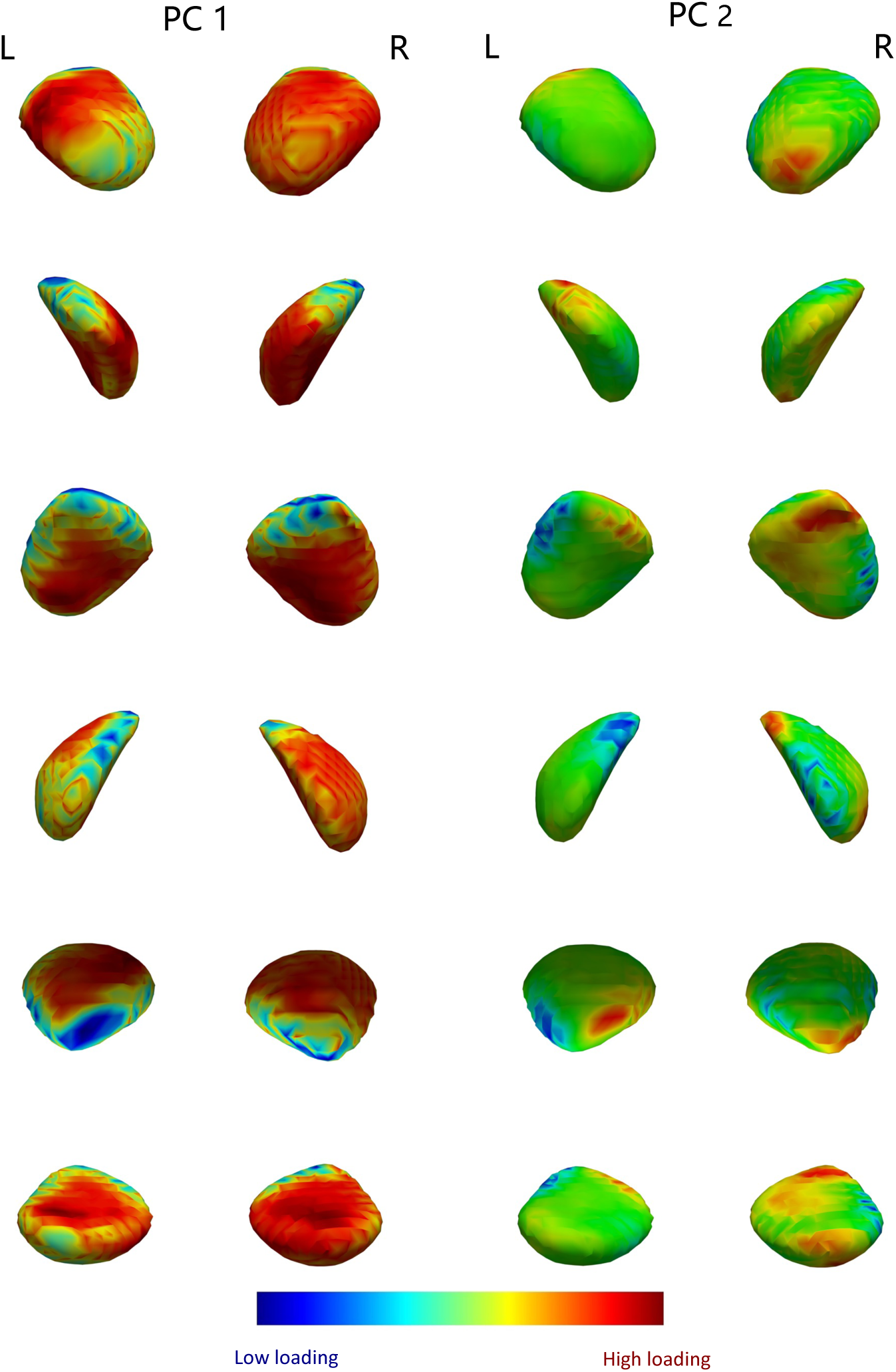
The two primary principal components of the time-varying relationship between amygdala shape and social anxiety from 7 to 16 years in the bilateral amygdala.

Our TVEM analysis also revealed significant links between the deformation of various surface areas of the amygdala and social anxiety over one year between 7-16 years (see Figure 4). However, these correlations were particularly observed in a very small region located at the basal nucleus of the amygdala (see Figure 3). The amygdala regions significantly and negatively associated with social anxiety were concentrated in the superficial subnuclei and the anterior and lateral subnuclei of the laterobasal subnuclei (Figure 4A). The amygdala regions significantly and positively associated with social anxiety were concentrated in the superficial subnuclei (Figure 4B). Although Figure 4 shows a map that illustrates the duration of significant relationships between amygdala shape and social anxiety, it does not differentiate temporal differences in these associations from 7 to 16 years. For example, an increase in a given vertex area can correlate with higher social anxiety scores in early childhood or exhibit positive correlations with social anxiety during late adolescence. These scenarios appear identical as shown in Figure 4. To further delineate the temporal dynamics of amygdala-anxiety associations, we modeled time-varying associations between amygdala morphology and social anxiety for 772 vertices across the left amygdala and 810 vertices across the right amygdala, and then collected unit-specific correlation estimates at 100 intervals between 7 and 16 years. Principal component analysis (PCA) of the correlation changes (772/810 vertices by 100 age-point measures of unit-area-anxiety correlation) identified three PCs that account for more than 80% and 87% of the variance in changes in the area-anxiety correlation at the vertex level for the left and right amygdala, respectively.

To jointly consider the dominant gradient of PC1 and PC2, we rendered both component loadings simultaneously in a color-coded two-dimensional space and plotted correlation trajectories for sets of vertices at all 4 extremes in the two-dimensional PC loading space (Figure 6). In the bilateral amygdala, the variation in PC1 loadings captured the distinction between regions with little discernible correlation across childhood and adolescence vs. those with a significant relationship across childhood and adolescence (that is, blue vs. red trajectories; turquoise vs. yellow trajectories; Figure 6). However, in the bilateral amygdala, between regions with low PC1 loadings, the variation in PC2 loadings captured the distinction between regions with positive associations in middle childhood vs. those with little discernible correlation in mid-childhood (yellow vs. red trajectories; Figure 6). In the bilateral amygdala, among regions with high PC1 loads, variation in PC2 loads had little discernible effect on the trajectory of change in correlation. They only captured the distinction between regions with little discernible correlation throughout childhood and adolescence versus those with slight negative associations in early adolescence (turquoise vs blue trajectories; Figure 6).

**Figure 6:**
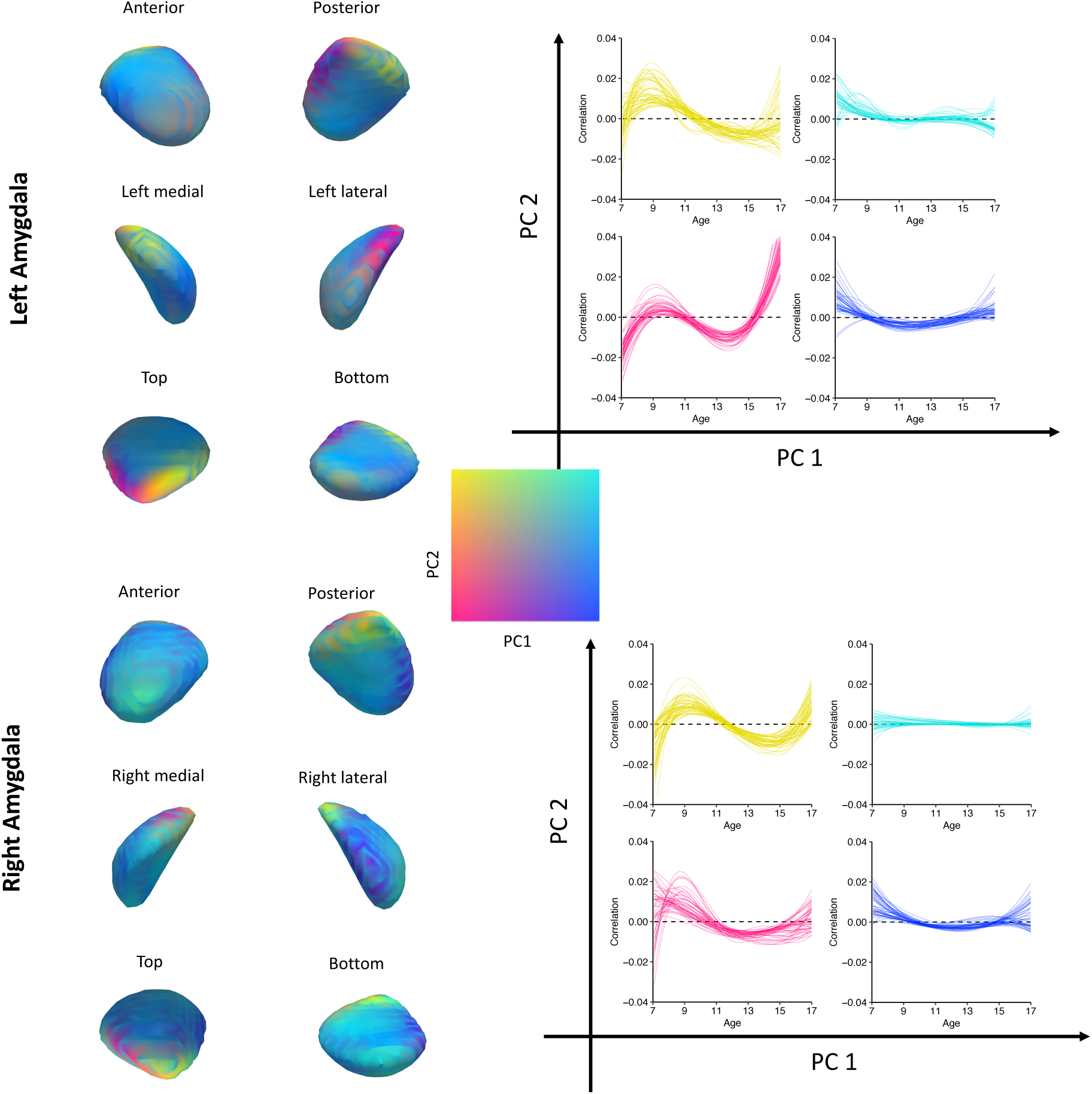
Combined visualization for both principal components of the time-varying relationship between amygdala shape change and anxiety from 7 to 16 years in the amygdala.

As shown in Figure 6, the superficial subnuclei of the bilateral amygdala (yellow) were where the regions of low PC1 and high PC2 loading were primarily concentrated, which tend to be positively related to social anxiety in the middle and late childhood and negatively related to social anxiety in the early and middle of adolescence. The low-loading regions of PC1 and PC2 were primarily concentrated in the laterobasal subnuclei of the bilateral amygdala (red), which showed negatively related social anxiety in early and middle adolescence. The regions of high PC1 and low PC2 loading were mainly concentrated in the laterobasal subnuclei of the right amygdala (blue), which showed negatively related social anxiety in early adolescence.

## Discussion

This study investigated the dynamic association between social anxiety and amygdala shape from childhood to adolescence, revealing a shift in the association between social anxiety and amygdala volume from positive to negative around the onset of puberty (10-11.5 years). Specifically, we found that the superficial nuclei of the bilateral amygdala initially exhibited a positive correlation with social anxiety in mid-childhood, transitioning to a negative correlation in early adolescence. Furthermore, a small region within the laterobasal nucleus showed a negative correlation with social anxiety at the beginning of puberty. These findings highlight a developmental shift in the link between the amygdala and social anxiety, influenced by distinct growth patterns in different regions of the amygdala.

Despite the lack of significant findings in the GAMM analysis, the TVEM analysis revealed age-specific correlations. The relationship evolved from a significant positive correlation in mid-childhood (ages 8-10) to a negative correlation in early adolescence (ages 11.5-13). The finding of a positive correlation in childhood is consistent with some previous pediatric studies (*21–23*), while the findings of a negative correlation in adolescence are consistent with existing adolescent (*24*) and adult (*28*) studies. Together, our findings reconcile inconsistencies in previous structural magnetic resonance imaging studies (*21–24*) and underscore the importance of considering the age range when examining the intricate relationship between amygdala volume and social anxiety.

The dynamic association between the amygdala and anxiety in childhood and adolescence suggests differential growth trajectories driven by social anxiety, that is, high social anxiety associated with amygdala overgrowth in childhood and delayed growth in adolescence. As depicted in Figure S2, the amygdala is initially larger in children with high social anxiety compared to their low-anxiety peers, but it does not show the age-related volumetric increase seen in the latter group. This pattern is consistent with findings on other disorders and stress-related changes. For example, initial episodes of major depression can lead to an increase in amygdala volume, which can decrease after recurrent episodes (*29*). Similarly, socioeconomic status impacts amygdala volume differently in developmental stages: lower socioeconomic status correlates with smaller amygdala volumes in adolescence but not in childhood (*30*).

The shifting association between social anxiety and amygdala volume may be explained through the lens of the allostatic load model (*31*), which posits that stress enhances cellular complexity in the amygdala, leading to increased dendritic branching and synaptogenesis. These changes result in increased excitation, surpassing inhibitory synapses growth, which can cause excitotoxic damage and cell death (*31*). Social anxiety specifically tilts the amygdala to a hyper-excitable state through multiple pathways including direct alterations in neurotransmitters like Glu and GABA, and indirectly through stress-induced changes in hormonal and neuropeptide signaling. Over time, this can lead the amygdala to a tipping point of over-excitation, resulting in structural diminution due to prolonged high stress levels (*31–33*). Second, the onset of puberty marks a critical period of increased sensitivity to interpersonal stress, often coinciding with a ”social reorientation” phase (*34,35*). Adolescents become increasingly sensitive to social signals and evaluations (*6*), which can amplify social anxiety and disrupt the normal excitation-inhibition balance within the amygdala, pushing it toward a hyperexcitable state. This state, if maintained from childhood, might culminate in reduced amygdala volumes (*31*). In contrast, the amygdala less affected during childhood and maintaining a low-load state could expand during this transition period. In summary, sensitivity to social stress during the transition from childhood to adolescence probably influences the volume of the amygdala, suggesting a change in the volume-anxiety association. The dynamic link between social anxiety and amygdala volume underscores the importance of early intervention and tailored approaches that recognize the evolving nature of the brain and its susceptibility to stressors in different stages of development. Future research should prioritize longitudinal studies to further explore these relationships and mechanisms, potentially informing preventative strategies and targeted therapeutic interventions for specific developmental periods.

Age-related associations between social anxiety and distinct geometrically deformed regions in the amygdala underpinned the findings of a shift in the association between social anxiety and amygdala volume around the onset of puberty. Most of the research informing neuropsychiatric research delineates the amygdala into three major subregions: the laterobasal, centromedial, and superficial subnuclei (*36,37*). The superficial subnuclei in the bilateral amygdala are positively related to social anxiety in childhood, revealing that the volume of the amygdala that increases with social anxiety in childhood has major expansion sites in the superficial region; the superficial and lateral laterobasal subnuclei in the bilateral amygdala are negatively related to social anxiety in adolescence, revealing that the volume of the amygdala that decreases with social anxiety in adolescence has major contraction sites. These findings emphasize the crucial roles these subnuclei play in the development of social anxiety and align with previous studies on the involvement of these subnuclei in social functioning (*38, 39*). Human neuroimaging studies have identified the laterobasal and superficial subregions of the left amygdala as key areas associated with perceived social support (*38*), and responsive to social stimuli (*39*). Structural anomalies of these subregions have been observed in individuals with autism, who often exhibit impaired social interactions (*40*). Comparative anatomical studies further support the evolutionary significance of these subregions; they have progressively expanded in primates (*41, 42*), especially in humans versus non-human primates (*41, 43*), suggesting a link to the evolution of social functionalities (*43*). Furthermore, studies in rodents showed that the deprivation of social relationships impaired neuronal structural plasticity and neurotransmission in the laterobasal (*44*) and superficial (*45*) subregions of the amygdala. Generally, our findings underscore the crucial roles of the superficial and laterobasal subnuclei in the developmental dynamics of social anxiety, providing significant insight into the neural underpinnings of social behavior.

The superficial subnuclei modulate the procession of olfactory and social information through specific functional connectivity with the limbic system (*36, 46*). Recent evidence has shown the roles of the superficial subnuclei subregions in the processing of both olfactory signals necessary for social recognition in mammals (*47, 48*) and facial expressions. The superficial subnuclei are believed to contribute significantly to social communication (*46, 49*), mediating social behaviors, including social recognition and rivalry between sexes, during typical development (*50, 51*). Therefore, the expansion of superficial subnuclei in childhood and the contraction in adolescence suggest that individuals who develop social anxiety in childhood and adolescence may exhibit abnormalities in processing emotional stimuli.

A portion of the laterobasal subnuclei in the bilateral amygdala showed negative associations with social anxiety in adolescence. The laterobasal subnuclei receive multisensory information from the prefrontal cortex (PFC), the hippocampus, the thalamus, and the visual and auditory cortices (*52,53*). Laterobasal subnuclei have been theorized to function as a ”gatekeeper” by assessing incoming sensory information and assigning emotional saliency to appropriate regions of the brain before the information is sent to other brain regions (*54*). Excessive contracting of the laterobasal subnuclei in adolescence can cause abnormal connections to multiple brain systems, such as decreased amygdala-medial prefrontal cortex connectivity, causing dysfunctional top-down control of information input (*22, 55*). Therefore, the contraction of the laterobasal subnuclei in adolescence suggests that individuals who develop social anxiety adolescence are more likely to exhibit abnormal emotion regulation.

Unlike the transition from positive to negative correlation with social anxiety experienced by the superficial nucleus, the significant negative correlation between the laterobasal nucleus and social anxiety only appears during adolescence. This result suggests that the susceptibility of the laterobasal nucleus and superficial nucleus to social stress differs at different age stages. These findings imply that early-onset and late-onset social anxiety may have different neural bases. Some researchers believe that early-onset anxiety may be caused by neurobiological vulnerabilities rather than environmental experiences, while the causes of anxiety symptoms that appear in adolescence may be more diverse (*22*).

Several limitations must be noted. First, the current study relied completely on self-report measures of social anxiety, which can be affected by false memory and misperception of participants. Future research utilizing multi-informant and observational paradigms with more powerful ecological validity and stable reliability to assess levels of social anxiety. Second, this study exclusively focused on social anxiety without examining its potential overlap with general social functions. Although prior psychological research suggests a mild association between social anxiety and the properties of social networks, it is not yet clear whether structural changes in the amygdala are specific to social anxiety or related to broader social functions. Future research should measure and compare the strength of these associations to better understand the neural correlates of social anxiety, which could provide critical insights for both theoretical and clinical applications. Third, our focus was primarily on the structure of the amygdala, thus neglecting other brain regions potentially linked to social anxiety. Given the extensive connections of the amygdala with other areas of the brain (*56*) and the potential link between the geometry and function of the human brain (*57*), future studies should explore broader in vivo imaging phenotypes, such as structural and functional connectivity of the amygdala with other regions of the brain. This could facilitate the exploration of more detailed aspects of amygdala growth in humans to more fully understand the neural mechanisms of social anxiety. Finally, although the current study had a large sample, future studies should include even larger clinical and non-clinical longitudinal samples to allow for a more nuanced investigation of the effects of various moderators. Furthermore, the macroscopic structural changes observed in the amygdala raise questions about the underlying cellular mechanisms (*58*). Future studies should investigate the cellular composition of the amygdala nuclei in children and adolescents to determine whether cell types and densities differ in relation to social anxiety. Evidence from postmortem studies has reported that cellular density and maturation in the various amygdala nuclei differ in children with social impairments compared to typically developing children and that these groups also differ in nucleic cellular loss and migration during childhood and adolescent development (*59, 60*). Currently, it is unknown which type of neurons were involved in these processes.

## Materials and methods

### Participants

The participants were recruited from the Chinese Color Nest Project (CCNP: https://ccnp.scidb.cn/en), an accelerated longitudinal initiative for studies of brain-mind development throughout human life (*61, 62*). Acceleration was implemented by combining cross-sectional and longitudinal designs to avoid long-term follow-up (*63,64*). The samples belong to the development component of CCNP (devCCNP), collected at Southwest University, Chongqing, China (devCCNP-CKG) (*65*). The devCCNP aims to chart normative brain development trajectories within the Chinese population throughout the school years. Eligible participants had no neurological or psychiatric conditions, did not use psychotropic drugs, and had estimated intelligence quotients of 80 or more. The devCCNP-CKG dataset included data from 201 participants aged 6-17 years who were invited to participate in three consecutive waves of data collection at intervals of approximately 1.25 years (*66, 67*). At these time points, T1-weighted MRI examinations were performed and the MRI images were visually inspected to exclude those with significant artifacts of head movement and structural abnormalities. After this initial quality control, the adopted samples included 427 scans from 198 participants (105 females and 93 males). The distribution of participants with one, two and three scans was, respectively, 48, 71, and 79, with an average of 2.16 scans per participant. The study received approval from the review committees of the participating institutions (Institute of Psychology at Chinese Academy of Sciences and Southwest University).

Social anxiety scores were measured with the Chinese version of the Social Anxiety Scales for Children (SASC) at the three time points (*68*). SASC is a ten-item self-report measure, designed for children aged 7-16, showing high reliability and validity in Chinese children samples (*69, 70*). The Cronbach’s α for the current sample was 0.79, 0.84, and 0.87, respectively, at the three assessments. After excluding participants who did not have social anxiety measurements, the study included 385 scans from 185 participants aged 7-16 (96 females and 89 males, as detailed in Table 1). The distribution of scans per participant was as follows: 52 participants (33 females; 19 males) with one scan, 66 (26 females; 40 males) with two scans, and 67 (37 females; 30 males) with three scans, averaging 2.08 scans/participant (see demographic details in Table 3).

**Table 3:** Sample characteristics and descriptive statistics.

|  | **Wave 1** | **Wave 2** | **Wave 3** |
| --- | --- | --- | --- |
| Number of participants | 166 | 132 | 87 |
| Female/Male | 90/76 | 62/70 | 44/43 |
| Average age (SD) | 11.66 (2.81) | 11.70 (2.30) | 12.25 (2.01) |
| Age range (years) | 7-16 | 7-16 | 9-16 |
| Average anxiety score (SD) | 6.02 (4.23) | 5.61 (4.55) | 5.00 (3.84) |
| Left volume (mean ± SD: cm<sup>3</sup>) | 1.46 ± 0.18 | 1.48 ± 0.18 | 1.48 ± 0.21 |
| Right volume (mean ± SD: cm<sup>3</sup>) | 1.46 ± 0.18 | 1.49 ± 0.18 | 1.50 ± 0.21 |

### MRI acquisition

All participants underwent MRI examinations with a Siemens Trio^TM^ 3.0 Tesla MRI scanner. A high-resolution magnetization-prepared rapid gradient-echo (MP-RAGE) T1 sequence (matrix = 256 × 256, FOV = 256 × 256 mm^2^, thickness of the slices = 1mm, repetition time (TR) = 2600 ms, echo time (TE) = 3.02 ms, inversion time (TI) = 900 ms, flip angle = 15*^◦^*, number of slices = 176) was obtained for each participant.

### Volumetric MRI preprocessing and segmentation

Imaging data were first anonymized using the **facemasking** toolkit (*71*), tailored for Chinese pediatric brain templates (*67*), which has been incorporated into the Connectome Computation System (*72*). The anonymized images were then processed through the online image processing system *volBrain* (http://volbrain.upv.es) for brain extraction (*73*). During this online computational procedure, the extracted individual brains were also cleaned by denoising using spatially adaptive non-local means and correction for intensity normalization in *volBrain*. All of these pre-processed brain volumes were in their native space and ready for subsequent manual and automatic tracing procedures. We verified that the impacts of face masking on brain extraction and preprocessing are trifling by visually inspecting each individual image.

Anatomically trained raters QZ (first author Quan Zhou) and ZQZ performed manual amygdala segmentation in native space using ITK-SNAP software (version 3.8.0) (*74*). The anatomical boundaries of amygdala structures were defined and segmented according to the protocol described in (*75*). This protocol has been demonstrated to achieve almost perfect intra-rater and inter-rater reliability. Reliability was quantified using the intraclass correlation coefficient (ICC), classified into levels of slight [0, 0.20), fair [0.20, 0.40), moderate [0.40, 0.60), substantial [0.60, 0.80), or almost perfect [0.80, 1] (*76, 77*). To assess the reliability of the implementation of the protocol, QZ and ZQZ independently traced the volumes of amygdala of 30 scans (from 30 individuals at baseline, balanced by age and sex) on two occasions, two weeks apart.

The manual tracing protocol exhibited almost perfect reliability for measuring amygdala volumes. Specifically, both intra-rater and inter-rater reliability was obtained for manually segmented amygdala volumes. The inter-rater ICCs were around 0.88 with 95%CI = [0.80, 0.96] for the left amygdala and 0.89 with 95%CI = [0.83, 0.95] for the right amygdala. The intra-rater ICCs were also almost perfect: 0.91 with 95%CI = [0.82, 0.96] for the rater ZQZ and 0.95 with 95%CI = [0.89, 0.97] for the rater QZ. These psychometric results confirmed that the raters’ manual tracings serving as the gold standard can be used as the reference for comparing against automatic segmentation methods.

### Shape analysis

The nonparametric statistical shape modeling (SSM) approach was used to reconstruct the template or atlas of the human amygdala, that is, the shape of the 3D anatomic mean amygdala of all subjects, without the need for additional manual landmarking (*78, 79*). Of note, this method simultaneously computes the mean shape from a 3D shape population and the deformation vectors that adjust this mean shape at the group level to each participant’s shape at the individual level. The mean and deformation vectors thus numerically describe all 3D head shape features of the population and allow statistical analyses of shape variations within a consistent framework. The steps of the statistical shape analysis pipeline are described in detail below.

#### Meshing and alignment

This step involves rigid transformations to align the samples for group modeling and analysis. The bilateral amygdala segmentation results, achieved by manual tracing, were refined using segmentation-based grooming tools and Python scripts developed by Cates et al. (*80*). During this computational neuroanatomy procedure, image padding, shape-based alignment (including center-of-mass and non-shrinking rigid alignment), and image cropping were implemented. Subsequently, surface meshes of aligned amygdala segmentations were generated with marching cubes on the binary masks using open-source VTK software (*81*).

#### Atlas construction and deformation capture

Following the recommendations in the *Deformetrica* guide (*78*), we employed *Deformetrica* (ver. 4.3) on a Linux operating system for our analyses, generating a statistical atlas of all shapes. This process involved identifying control points in areas of high variability between participants and calculating momenta for these points, which represent directional shape variations of the atlas. We established point-to-point correspondences for each shape, identifying 772 control points for the left amygdala and 810 for the right amygdala. Numerical estimates of surface area at each vertex were derived by allocating one-third of the adjoining triangle’s surface area to each vertex and summing these for all associated triangles. These estimates of vertex-level surface area at 772 and 810 control points for the left and right amygdala, respectively, allowed us to spatially analyze amygdala shape variations during school-age development.

### Statistical analysis

#### Associations between amygdala volume and social anxiety

We used the generalized additive mixed model (GAMM) to investigate the relationship between social anxiety and amygdala volume. GAMM is a semiparametric regression approach and allows flexible and efficient exploration of potentially nonlinear relationships between social anxiety and amygdala volume without the need for predefined assumptions about their interaction (*82–84*). They are particularly advantageous for analyzing data from repeated measurements, such as our accelerated longitudinal samples of developing brains, by accommodating both within-subject dependencies and developmental variabilities among participants (*85, 86*). We implemented GAMMs using the `mgcv` package in R (*87*), following the specified formula:

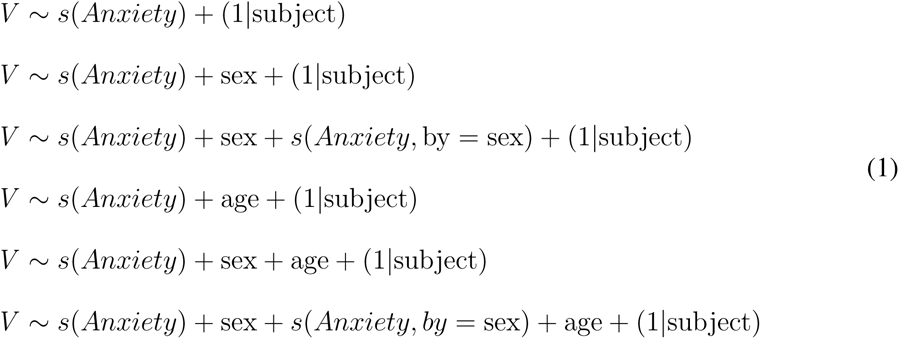

The first GAMM treats the volume of the amygdala, denoted *V*, as a smoothing function of social anxiety. The second GAMM introduces sex as a fixed-effect factor on the basis of the first model, while the third GAMM further includes an anxiety-sex interaction to evaluate sex-based differences in response slopes. To exclude age-related volume changes, subsequent models (fourth to sixth) incorporate age as a fixed effect factor into the first three GAMMs, respectively. The fit of the model was assessed using the Akaike Information Criterion (AIC). Each model was tested against a null anxiety effect model. The model with the lowest AIC and a significant deviation from the null was selected as optimal. Analyses and visualization were performed using R packages `mgcv` (*83*) and `ggplot2` (*88*).

#### Timing-varying relationships between amygdala volume and social anxiety

We explore the evolving relationship between amygdala volume and social anxiety from childhood to adolescence using the time-varying effect model (TVEM, (*89*)) using the % TVEM SAS macro version 3.1.1 (*90*). TVEM extends traditional multilevel models by allowing the association between variables to vary over time without preset functional forms (*89*). In our TVEMs, both the intercept and anxiety’s impact were modeled as time dependent, with 10 knots for each and sex considered as a constant factor. We employed the P-spline method to fit the model, ensuring sufficient complexity in the shape of the curve. The P-spline method automatically determines the degree of complexity of each coefficient function in the macro. Furthermore, it applies robust standard errors automatically to address the nonindependence of repeated assessments, a feature intrinsic to all longitudinal data, thereby enhancing the method’s validity in analyzing such data.

#### Associations between amygdala shape and social anxiety

We introduced a series of GAMMs to investigate the potential associations between the amygdala shape and social anxiety scores:

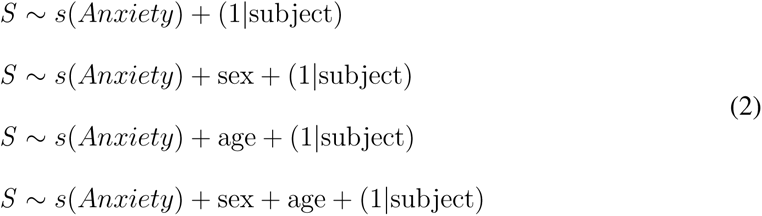

The first GAMM treats the surface area of the amygdala, denoted as *S*, as a smoothing function of social anxiety. The second and third GAMMs introduce sex and age, respectively, as fixed-effect factors on the basis of the first model to assess the sex and age differences at the intercept of the regression curve. The fourth model included both sex and age as fixed-effect factors. AIC was used to evaluate the fit of the models. Beyond direct estimation of the link between amygdala shape and social anxiety via GAMM, TVEM was also applied to explore the evolving relationship between the area of individual surface vertices and social anxiety in the age range. In our TVEMs, both the intercept and anxiety’s impact were modeled as time dependent, with 10 knots for each and sex considered as a constant factor. The P-spline method was used in all analytical procedures.

To visualize regional changes in the patterns of time-varying correlation trajectories in the bilateral amygdala, we employed two methods. First, to highlight regional differences in the significance and duration of the correlation between amygdala deformation and social anxiety from ages 7 to 16, vertex-wise maps were generated. These maps succinctly depict the interregional differences in significant correlations at the vertex level on the surface of the amygdala. Additionally, for a more detailed analysis, we calculated the area-anxiety correlation values for 100 age points evenly distributed throughout the age range for each surface vertex. Subsequently, the principal component analysis (PCA) was performed on these correlation matrices (with vertices as rows, ages as columns, and correlation estimates as values). The principal components (PCs) of correlation trajectories could then be visualized on the surface of the amygdala to display the primary spatial patterns of variation in the temporal dynamics of the shape of the amygdala and anxiety associations. Two PCs were identified as optimal on the basis of the scree plot analysis. We produced separate and combined surface maps, color-coding each vertex according to its PCA loading in the two-dimensional PC space. To explore the variations in correlation trajectories indicated by the two PCs, we modeled and visualized age-related associations for vertices at the extreme ends of the PC loading space. This analysis encompassed four distinct scenarios within the two-dimensional space: high PC1 & low PC2, low PC1 & high PC2, high PC1 & high PC2, and low PC1 & low PC2. We identified each scenario by selecting the top 10% of the vertices according to their combined scores on PC1 and PC2 loadings. For example, to isolate vertices at the extreme for high PC1 & low PC2, we calculated PC1 loading × (−1×PC2 loading) for each vertex and took the top 10% from a ranking list.

## Acknowledgments

Chinese Color Nest Consortium (CCNC) receives funding support from the Major Fund for International Collaboration of the National Natural Science Foundation of China (81220108014), the Key Research Program of the Chinese Academy of Sciences (KSZD-EW-TZ-002), the Beijing Municipal Science and Technology Commission (Z161100002616023) and the National Basic Science Data Center ’Interdisciplinary Brain Database for In vivo Population Imaging (ID-BRAIN). CCNC wishes to thank all community partners, research participants, and families. Xi-Nian Zuo is partly supported by the STI 2030 - the major projects of the Brain Science and Brain-Inspired Intelligence Technology (2021ZD0200500) and the Start-up Funds for Leading Talents at Beijing Normal University. Quan Zhou and Yin-Shan Wang are supported by the China Postdoctoral Science Foundation (2023M740301 and 2022M710432).

## Supplementary Materials

### Time-varying relationships between age, sex, and amygdala volume

We determined the effects of age on individual differences in bilateral amygdala volume (the time-varying intercept), with sex included as a time-invarying covariate. The best fit of the model was identified for the intercept in the volume of the left and right amygdala: the bilateral amygdala exhibited inverted U-shaped growth patterns: robust volume increase during childhood and early adolescence, and gradual decrease in middle adolescence in the bilateral amygdala (the peak age around 14 years old) (Supplemental Figure S1).

**Figure S1:**
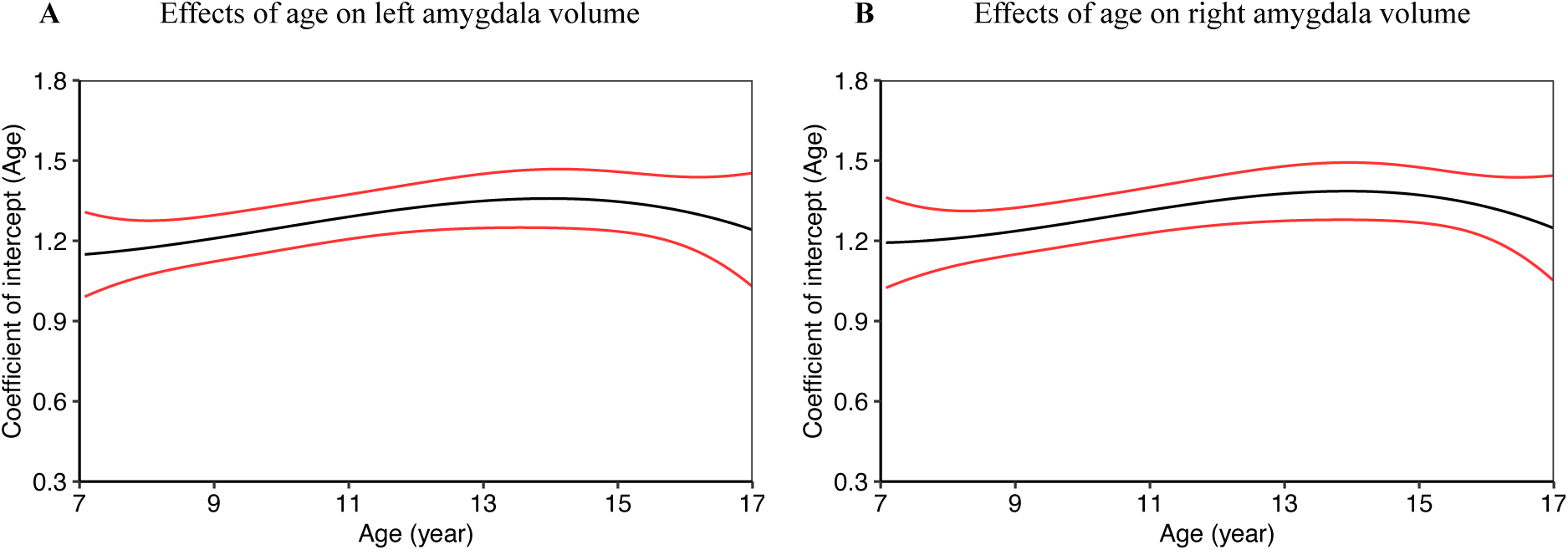
Time varying effects of age (intercept) on A) left and B) right amygdala volume.

### The growth curve of social anxiety in childhood and adolescence

We use the following GAMM to examine age-related changes of social anxiety by including social anxiety as a smoothing function of age. As shown in Supplemental Figure S2, GAMM revealed the growth patterns of social anxiety: linearly increasing with age throughout childhood and adolescence (edf=1, *F*=63.18, *p*<0.001).

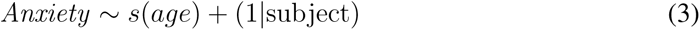

**Figure S2:**
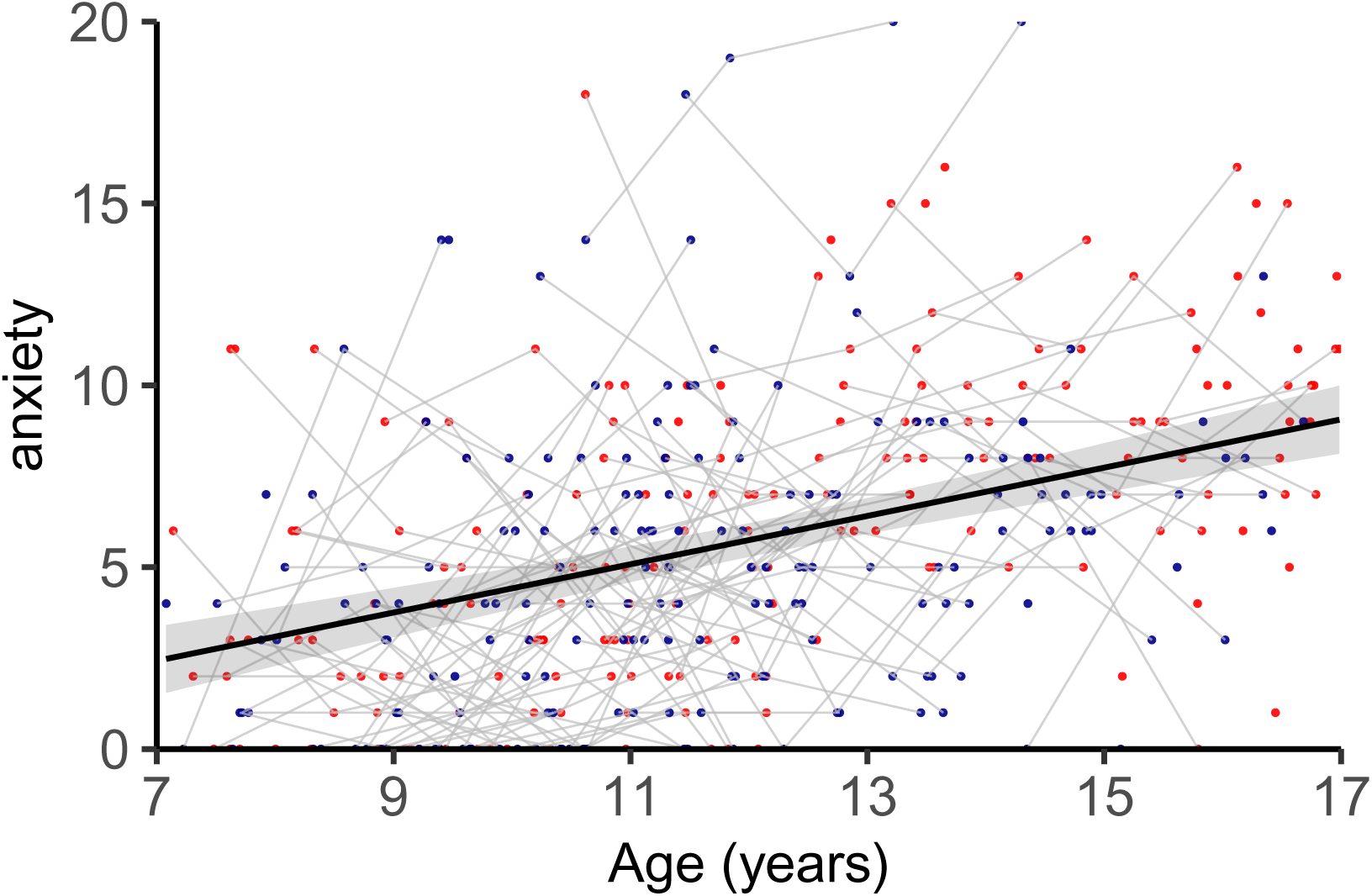
Growth curve of social anxiety in childhood and adolescence.

### Growth curves of the amygdala volume for high and low levels of social anxiety

To explore the growth curves of the amygdala volume for high and low levels of social anxiety, we used the following GAMM to examine the developmental changes of the amygdala volume by including volume as a smoothing function of age. The groups were: high level of the social anxiety group (range of anxiety value 8-20) and low level of the social anxiety group (range of anxiety value 0-3). As shown in Figure S3, the GAMM revealed the growth patterns of the volume for bilateral amygdala of two groups: non-linearly increases for bilateral amygdala volume of low anxiety group (left amygdala: *F*=4.11, *p*=0.016; right amygdala: *F*=4.70, *p*=0.012); no significant change with age for bilateral amygdala volume of high anxiety group (left amygdala: *F*=0.28, *p*=0.596; right amygdala: *F*=0.08, *p*=0.779).

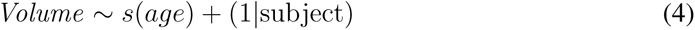

**Figure S3:**
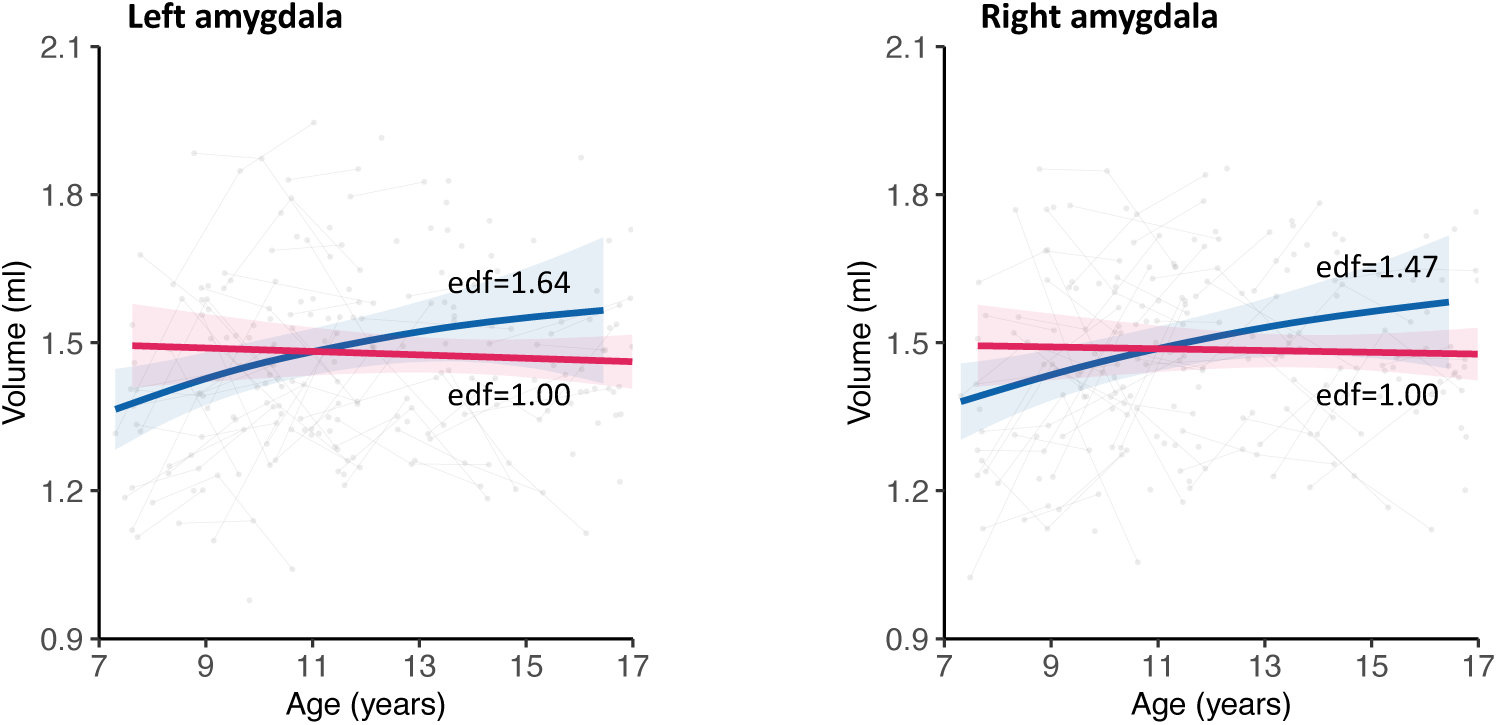
Growth curves of amygdala volume for high and low levels of social anxiety. The blue color indicates curves for the low social anxiety group, while the red color for the high social anxiety group’s curves. The curves are surrounded by shaded 95% confidence intervals. The low anxiety group developed along non-linearly increasing patterns for bilateral amygdala (ps<0.01), while the high anxiety group had no significant changes with age.

### The growth associations of the amygdala shape with social anxiety throughout childhood and adolescence

We introduce two GAMMs as follows to further validate the geometric deformation specific to age stage associated with social anxiety in mid-childhood and early and middle adolescence shown in Figure 6.

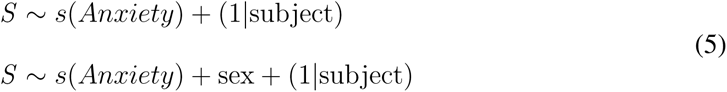

The first GAMM treats the surface area of the amygdala, denoted as S, as a smoothing function of social anxiety. The second GAMMs introduce sex as a fixed effect factor on the basis of the first model to assess the sex differences at the intercept of the regression curve. AIC was used to evaluate the fit of the models. Regarding the post hoc nature of this procedure and the robustness consideration (27), vertex-wise statistical significance of the association tests remained uncorrected for multiple comparisons. As shown in Figure S4, the distribution map of the p-values revealed areas at the vertex level of the age stage (hot spots) of the development of the anxiety-related areal, consistent with Figure 6 and Figure 4. For the left amygdala, the geometric deformation related to social anxiety in mid-childhood (8-11 years) has major expansion sites in the superficial region; the geometric deformation related to social anxiety in early and middle adolescence (12.5-15 years) has major contraction sites in the superficial and basolateral region. For the right amygdala, the geometric deformation related to social anxiety in middle childhood (8-11 years) has major expansion sites in the superficial region; the geometric deformation related to social anxiety in early and middle adolescence (11.5-16 years) has major contraction sites in the superficial and basolateral region.

**Figure S4:**
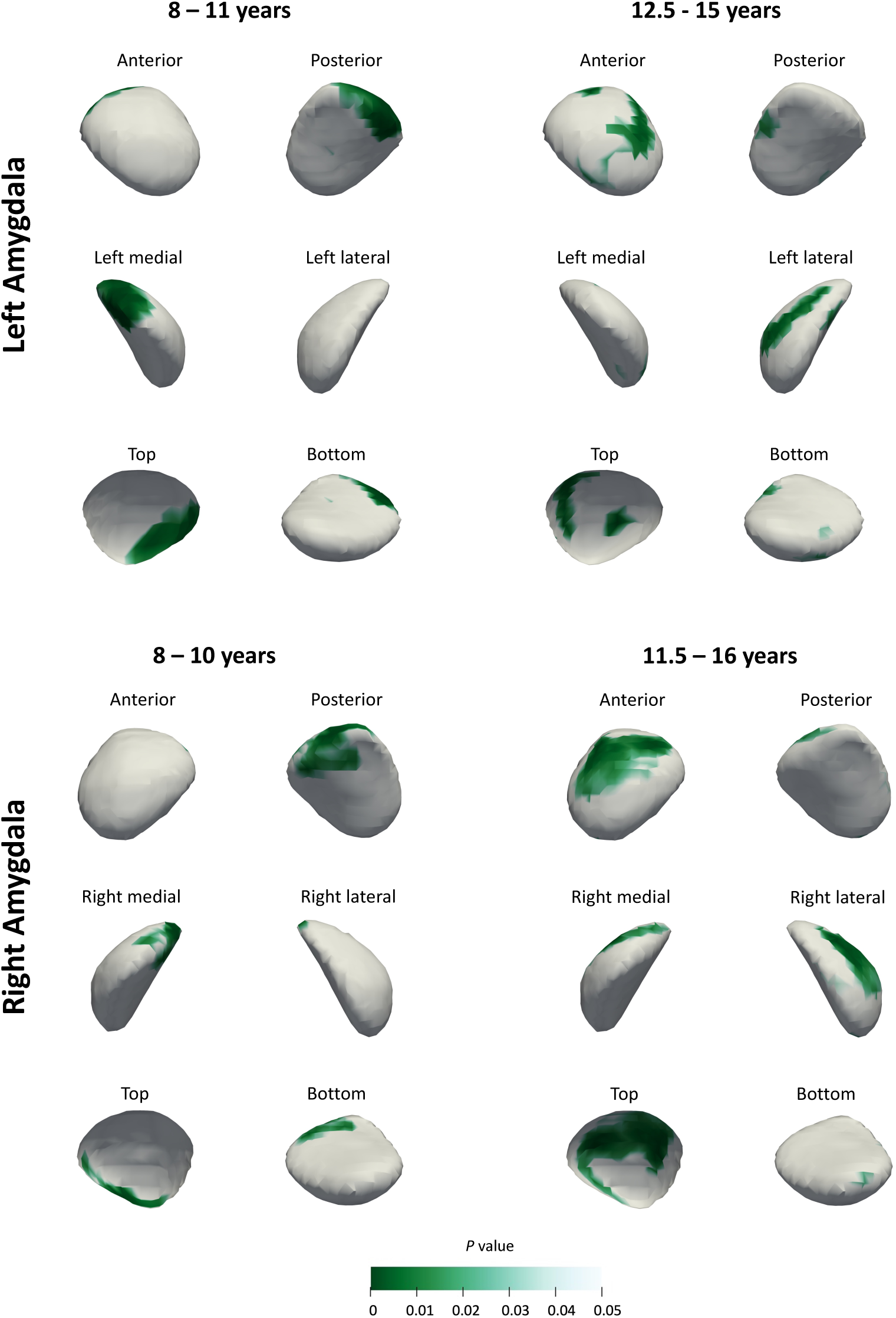
Age stage-specific statistically significant surface deformations associated with social anxiety. The anterior, posterior, medial, lateral, top, and bottom view of P-statistic maps were shown. The significant surface vertices (*p* < 0.05) are colored in green.

